# Structure-Function Dynamics in Healthy Cognitive Aging: A Graph Signal Processing Approach

**DOI:** 10.1101/2025.09.23.678034

**Authors:** Clément Guichet, Sophie Achard, Martial Mermillod, Monica Baciu

## Abstract

Network neuroscience has significantly advanced our understanding of how structural and functional connectivity evolve during healthy neurocognitive aging. Yet, integrative studies linking structural and functional brain organization with cognitive performance remain relatively limited. In this study, we analyzed resting-state functional MRI and diffusion-weighted imaging data from 600 healthy adults aged 18 to 88, drawn from the CamCAN dataset. Using a graph signal processing framework, we investigated how structural connectivity constrains functional brain signals across the adult lifespan. Our results reveal that control and semantic cognitive systems exhibit distinct age-related patterns of structure-function reorganization, potentially reflecting a shift in integrative processing during midlife. Notably, our findings suggest that structurally-coupled sensorimotor integration plays a crucial role for regulating these systems. Until midlife, it accompanies structurally-decoupled activity in transmodal cortices to sustain cognitive control. In parallel, it likely supports the formation of embodied internal models that leverage more structurally-decoupled semantic processes, thus contributing to maintain lexical production skills for longer in older adulthood. Taken together, our study offer new multimodal insights into how sensory-driven processes help reconfigure the healthy aging brain to support controlled semantic cognition.

## 1. Introduction

Neurocognitive aging is generally presented as a gradual decline in control abilities such as attentional control, inhibitory control, as well as episodic and working memory processes (Salthouse, 2019). However, certain cognitive functions, such as vocabulary and semantic knowledge, remain stable or even improve with age (Kavé & Halamish, 2015; Shafto et al., 2017). More broadly, this increased “semanticization” (Spreng et al., 2018) is thought to reflect a shift towards a distinct “neurocognitive mode” in older adulthood, marked by a greater reliance on accumulated knowledge and past experiences (Spreng & Turner, 2021).

While previous research has explored candidate structural and functional substrates separately, a comprehensive analysis of the age-related structural constraints shaping functional activity remains insufficient. This is especially relevant as upcoming models are geared towards multimodal integration between cognitive, structural, and functional imaging modalities (Avena-Koenigsberger et al., 2018; Honey et al., 2010; Lynn & Bassett, 2019). Relatedly, extensive research supports that the structure-function architecture as a whole is a fundamental property of brain organization with high predictive power for individual-level cognition (Feng et al., 2024) (Dong et al., 2024; Griffa et al., 2022; Gu et al., 2021; Zimmermann et al., 2018). For example, a growing body of word describe an architecture following the unimodal-to-transmodal gradient of connectivity (Margulies et al., 2016) : stronger structure-function coupling in primary sensory and motor cortices at one end of the hierarchy, and greater decoupling in transmodal association cortices at the other end (Baum et al., 2020; Fotiadis et al., 2023; Suárez et al., 2020; Sun et al., 2022; Vázquez-Rodríguez et al., 2019; Wang et al., 2019; Yang et al., 2023).

With this in mind, our current study aims at exploring the structure-function coupling and decoupling dynamics that accompany both declining and preserved cognitive abilities in healthy aging (Loaiza, 2024). To characterize the interaction between declining control and preserved semantic trajectories with age, language offers a convenient framework (Hagoort, 2023; Pinker, 2008).

Indeed, while language comprehension remains intact until very old age, language production often becomes more challenging beyond midlife as it relies on control processes for lexico-semantic access and retrieval (Baciu et al., 2021; Oosterhuis et al., 2023; Verhaegen & Poncelet, 2013). Specifically, lexico-semantic selection in older adulthood is thought to be disrupted by intrusive thoughts due to reduced inhibitory control (Baciu et al., 2016; Blanco et al., 2016; Hoffman & Morcom, 2018). This disruption manifests behaviorally in more frequent tip-of-the-tongue situations and longer picture naming latencies, particularly for words with limited semantic connections (Benítez-Burraco & Ivanova, 2023). In sum, these findings join previous works showing that control deficits compromise semantic access in older adulthood, but that a densely connected repertoire of learned semantic associations may help individuals mitigate these deficits – a “semantic strategy” – which maintains word access and retrieval for longer (Gollan & Goldrick, 2019; Krethlow et al., 2024).

At the brain level, the dual trajectory of control and semantic neurocognitive systems with age have been formalized in models such as LARA (*Lexical Access and Retrieval in Aging*) (Baciu et al., 2021; Baciu & Roger, 2024), and more recently within a connectomic framework via SENECA (*Synergistic, Economical, Nonlinear, Emergent, Cognitive Aging*) (Guichet, Banjac, et al., 2024). These models acknowledge midlife as a critical transition period (Beck et al., 2021; Dohm-Hansen et al., 2024; Hennessee et al., 2022; Roger et al., 2023; Shafto & Tyler, 2014), putting emphasis on age-specific strategies used to uphold lexical production – control-based in younger adults and semantic-based in older adults.

More specifically, in younger adults, flexible modulation of functional connectivity between the Default Mode (DMN) and Fronto-Parietal (FPN) networks has been shown to sustain goal-directed semantic access and retrieval internal (Chiou et al., 2023; Jackson, 2021; Menon, 2023). thereby suggesting a direct link with lexical production (Guichet, Banjac, et al., 2024). This strategy particularly involves medio-parietal connectivity within the posterior cingulate cortex, which serves as a key DMN-FPN interface (Fan et al., 2019; Leech & Sharp, 2014) during semantic control tasks (Krieger-Redwood et al., 2016). Structurally, younger adults also benefit from optimal microstructural integrity along subcortico-frontal and dorsal association pathways (Bennett et al., 2010; Boban et al., 2022; Bonifazi et al., 2018; Troutman et al., 2022; Zahr et al., 2009), the latter being associated with inhibition and cognitive flexibility (Ribeiro et al., 2023; Rizio & Diaz, 2016; Troutman et al., 2022). Beyond midlife, more rigid DMN-FPN functional connectivity as described by the DECHA model (*Default-Executive Coupling Hypothesis in Aging*; *Spreng* & Turner, 2019) and loss of white matter microstructural integrity in the associated pathways may lead to poorer filtering of task-irrelevant thoughts that compromised controlled semantic access.

In parallel, the expansion of the semantic space (Cutler et al., 2025) is thought to peak around midlife and remain stable until late older adulthood. Semantic hubs are thought to remain largely preserved among older adults exhibiting successful aging (Garcia et al., 2022, 2024). Structurally, previous work has associated this expansion with ventral pathways such as the IFOF and ILF (Baciu et al., 2021; Giampiccolo et al., 2025), as well as a left-dominant subcortico-sensory network of white matter fibers (Guichet, Roger, et al., 2024) to maintain lexico-semantic representations (Crosson, 2021). Functionally, further work also highlighted more dynamic interactions within sensorimotor networks (Guichet, Banjac, et al., 2024). Although the role of sensorimotor processing in this strategy remains unclear, some have emphasized that it may enhance the comparison of sensory information to long-term semantic memory traces (Brown et al., 2022).

Overall, the LARA and SENECA models illustrate that lexical production provides a valuable framework for understanding a multimodal shift in midlife that bridges structure, function, and cognition, specifically targeting how older adults regulate the flow of information between semantic, sensory and control systems beyond a left-lateralized core language network (Forkel & Hagoort, 2024; Hagoort, 2023; Hertrich et al., 2020; Roger et al., 2022; Thiebaut de Schotten & Forkel, 2022).

### The present study

Our main hypothesis is that midlife is a critical period at the junction between two structure-function architectures. (1) The first architecture related to cognitive control. It may be driven by the decoupling of the DMN and FPN functional activity from underlying structural constraints, offering flexible attentional processes for controlled access to the semantic store. This hypothesis is based on previous work showing a relationship between higher decoupling and sustained attentional performance, verbal learning and retrieval (Griffa et al., 2022), as well as with functional diversity and cognitive flexibility (Fotiadis et al., 2024; Wu et al., 2020; Yeo et al., 2015). We expect this architecture to show significant decline beyond midlife, particularly affecting medio-parietal areas. (2) The second architecture relates to semantic access. It could be regulated by enhanced coupling of sensorimotor cortices, providing the foundation for enhanced lexico-semantic access from midlife onwards. In line with this hypothesis, high coupling in sensorimotor areas have been proposed to reflect fast and reliable interfacing with the environment (Preti & Van De Ville, 2019).

To examine these hypotheses, we propose to combine the information carried by two imaging modalities – diffusion-weighted imaging (DWI) and functional MRI – in a population-based sampled of healthy adults aged 18-88 (Cam-CAN et al., 2014). We adopt a graph signal processing approach (Abramian et al., 2021; Fotiadis et al., 2024; Lioi et al., 2021) that decomposes the structural connectome into harmonics – a graph spectral representation that capture modes of spatial variations – and expresses the fMRI BOLD signal as a function of this representation. The key idea is that functional activity driven by low-frequency harmonics is structurally coupled whereas activity driven by high-frequency harmonics is decoupled (Preti & Van De Ville, 2019).

## 2. Results

### 2.1. Comparison of structural connectome harmonics between subjects

Figure 1A illustrates the first harmonics representing modes of structural variations with increasing spatial frequency. The first harmonic captures spatial uniformity, while the second and third harmonics resolve left-right and anterior-posterior axes of connectivity, respectively.

**Figure 1.**
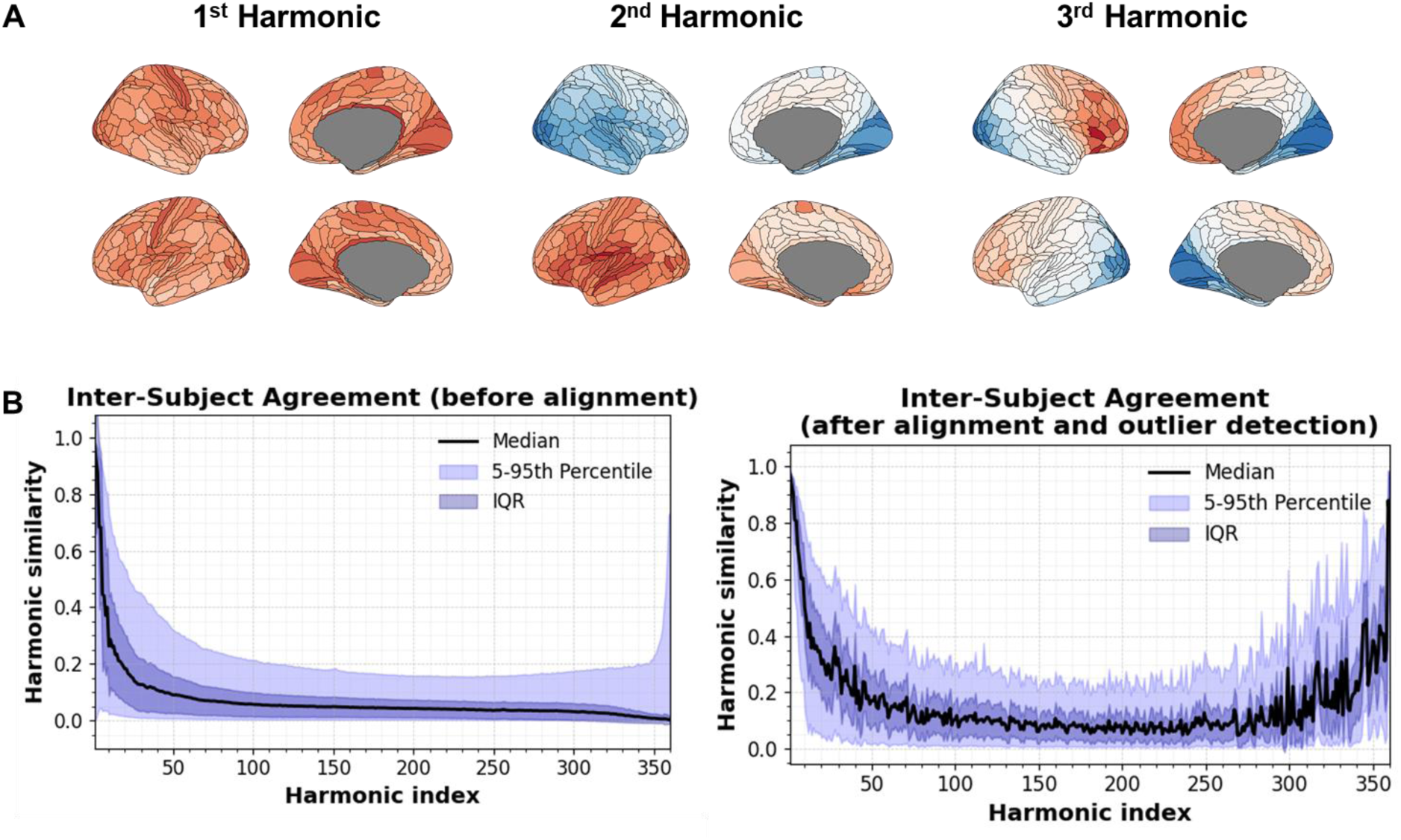
Harmonic inter-subject agreement. **(A) Group-averaged structural connectome harmonics**. The structural harmonics. Brain illustrations made with ggseg R-package (Mowinckel & Vidal-Piñeiro, 2020) **(B) Harmonic inter-subject agreement**. The right panel shows significant improvement in the agreement across all combinations of subjects. *IQR: inter-quartile range*

Figure 1B shows that subject-specific harmonics are naturally misaligned except for the ones capturing the main axes of structural connectivity. Following our alignment procedure, we observed significant improvement in the median inter-subject agreement (*p_wilcoxon_* < .001), enabling more reliable comparisons between subjects.

### 2.2. Age-related changes in the energy spectrum

To assess changes in structure-function relationship across the lifespan, we expressed the fMRI signal as a combination of the structural harmonics and analyzed the resulting energy spectrum. This spectrum indicates the contribution of each harmonic to the fMRI signal.

We observed a substantial increase in harmonic diversity indicating a broadening of the structural repertoire used to compose fMRI activity with age (*F* = 79.8, *p* < .001, *edf* = 1.4, partial R² = .17). To pinpoint the harmonics driving this effect, we binned the energy spectrum in 36 bins (one bin for 10 harmonics) and conducted the same analysis. We found that our effect is driven by the first 10 harmonics with a clear inflection point at age 54 (*F* = 22.33, *p* < .001, *edf* = 1.93, partial R² = .07), thus suggesting a broader recruitment of the main axes of structural connectivity in older adulthood.

To gain in interpretability, we projected the information carried by these 10 modes back in the spatial domain, resulting in structure-informed functional timeseries upon which further statistical analysis was performed. First, we derived integration/segregation graph metrics to confirm that these harmonics support functional integration (Figure 2B). Next, we examined how this integration is expressed spatially before and after the inflection point at age 54. Age-related differences occurred primarily along the medial wall, with older adults engaging more dorsal, anterior cingulate and posterior parietal regions while younger adults mostly engaged medio-orbito-frontal and insular regions (Figure 2A).

**Figure 2.**
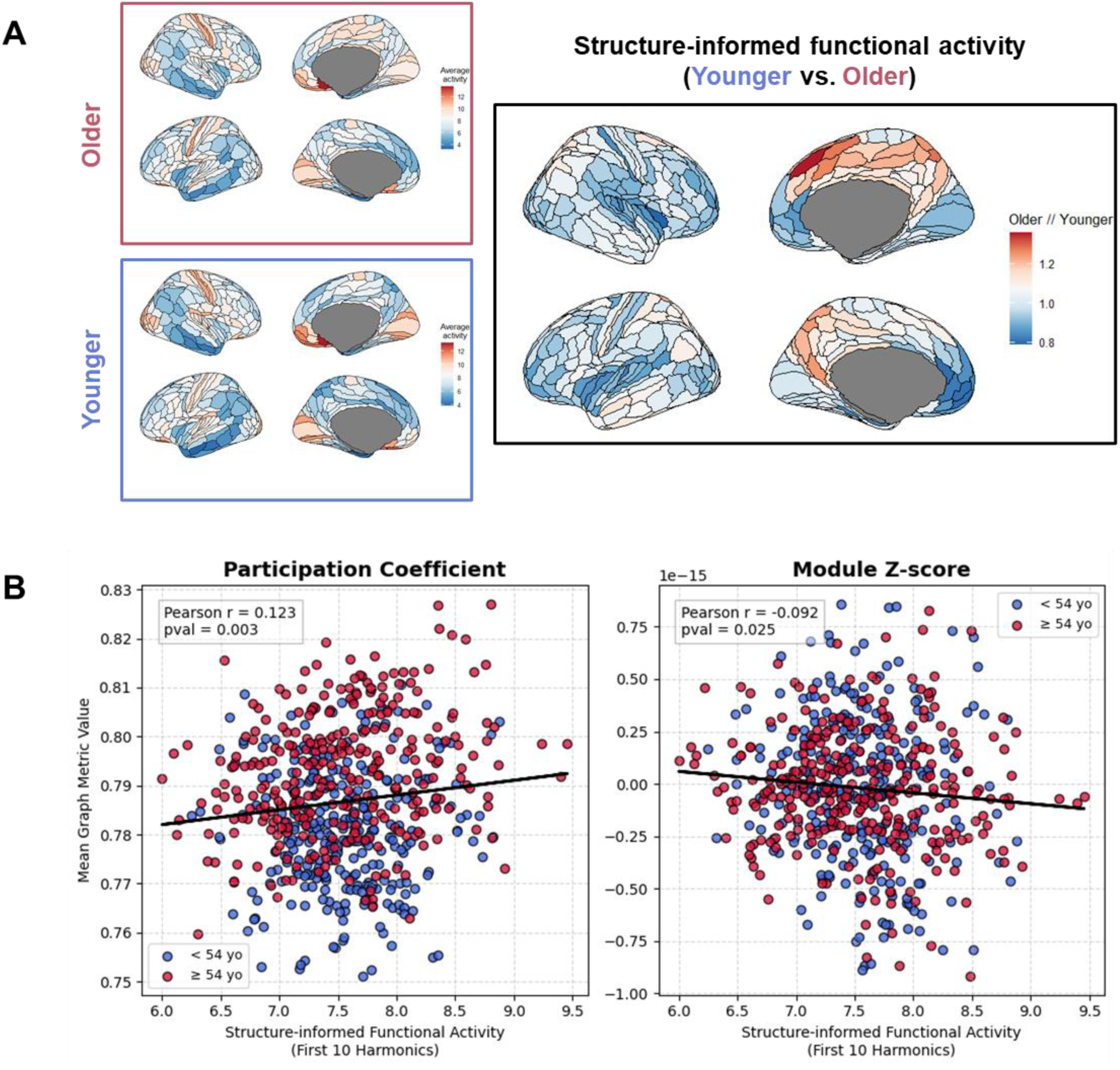
Structure-informed functional integration. **(A) Age-related effect**. Subject-specific maps were obtained by projecting the first 10 harmonic projected back into the space domain and taking the Euclidean norm across time. The age contrast illustrated on the right panel is the ratio between the average older and average younger maps (± 54 yo) as shown on the left panels. Brain illustrations made with ggseg R-package (Mowinckel & Vidal-Piñeiro, 2020) **(B) Correlation with graph metrics.** The participation coefficient is a measure of intermodular connectivity (integration); the module degree z-score is a measure of intra-modular connectivity (segregation). Computations made with the *bct* toolbox https://github.com/aestrivex/bctpy

We also found a small but significant reduction in temporal diversity beyond midlife (*F* = 6.2, *p* < .001, *edf* = 1.7, partial R² = −.02), meaning that the structure-function relationship becomes less specific in time.

### 2.3. Age-related changes in the structure-function decoupling index (SDI)

Next, we examined the information carried by low vs. high-frequency harmonics, respectively reflecting coupling vs. decoupling to structural constraints of each region. At the whole-brain level, healthy aging was associated with enhanced decoupling (*F* = 16.19, *p* < .001, *edf* = 1, partial R² = .03), suggesting reduced support from large-scale structural axes.

At the region level, and in line with our previous results, Figure 3A illustrates linear but also nonlinear alterations with an inflection point at age 54. Further analysis showed increased coupling with age in transmodal cortices such as the DMN and attentional networks such as the DAN and FPN. In comparison, increased decoupling with age was observed in the SMN, Visual, and Auditory networks.

**Figure 3.**
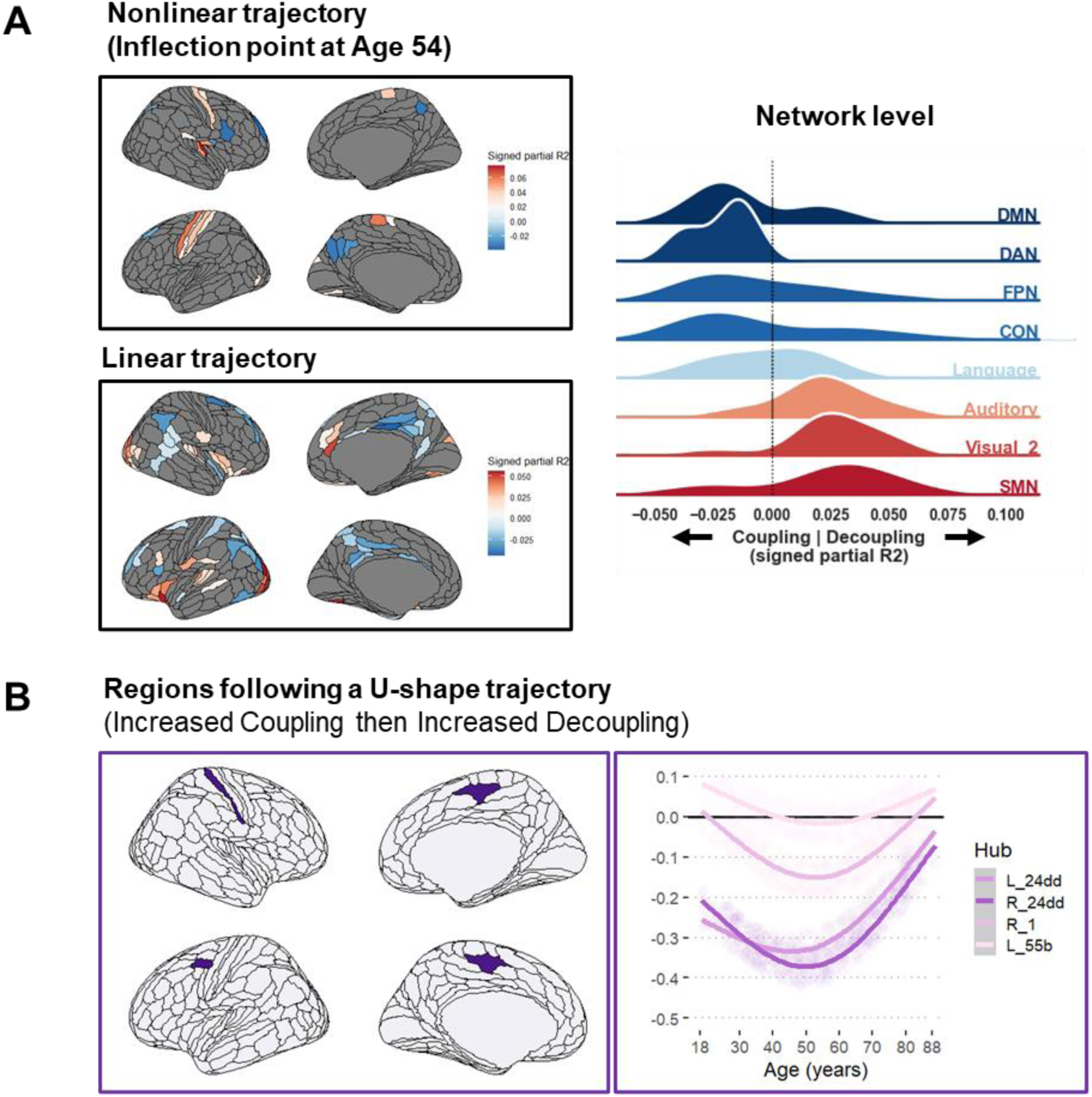
Structure-Decoupling Index (SDI). **(A & B) Age-related changes in SDI**. Reds indicates more decoupling; Blue indicates more coupling. Brain illustrations made with ggseg R-package (Mowinckel & Vidal-Piñeiro, 2020)

Moreover, Figure 3B illustrates key regions exhibiting a U-shape trajectory with age (i.e., greater coupling until midlife followed by greater decoupling), potentially acting as key structure-function interfaces during healthy aging. Further analysis based on white matter bundle segmentations ((https://github.com/MIC-DKFZ/TractSeg; Wasserthal et al., 2018) revealed that these regions straddle associative (e.g., 91% of fibers of left SLF II transit via these regions), callosal (CC4: 79% and CC7: 84%), and projection fibers (left optic radiations: 85%, left thalamo and striato-thalamo occipital pathways: 79% & 76%).

### 2.4. Structure-Function-Cognition analysis

We further sought to relate these changes to lexical production performance using the Partial Least Squares (PLS) correlation analysis.

In line with our hypothesis, we identified one major cognitive control component (LC1) which explained 72.6% (*p_FDR_* = .001) of the total shared variance and one semantic component (LC2) which explained 9.4% more variance (*p_FDR_* = .002). As shown in Figure 4, key lexical production tasks such as picture naming (BSR = 21.55/11.05) and verbal fluency tasks (BSR = 9.87/16.45) were positively correlated to both components, thereby confirming the cognitive control and semantic aspects of age-related lexical production when fusing imaging modalities.

**Figure 4.**
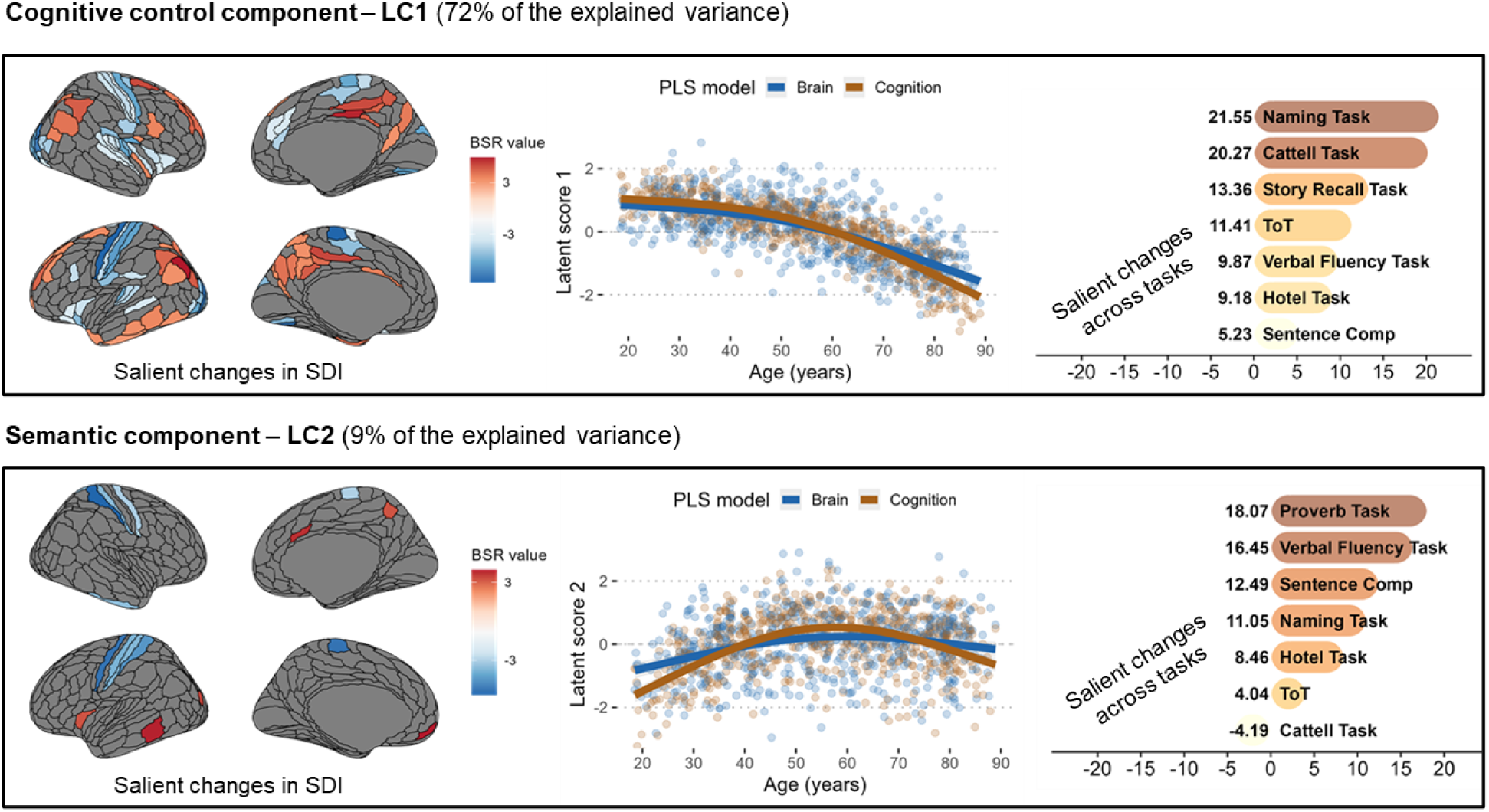
Structure-function-cognition analysis. The middle panels illustrate the trajectory of the first and second latent components (LC1 and LC2) inferred by the PLS. The left and right panels respectively report the brain and cognitive profiles correlated with each trajectory. Only bootstrap sampling ratios (BSR) ± 3 denote a robust contribution to the covariance patterns. Brain illustrations made with ggseg R-package (Mowinckel & Vidal-Piñeiro, 2020)

#### 2.4.1. Cognitive control – Latent component 1

At a cognitive level, LC1 was mainly associated with better picture naming performances (21.55), enhanced fluid abilities Cattell task (20.27), long-term memory (13.36) and fewer tip-of-the-tongue occurrences (11.41). These tasks performances significantly decreased beyond midlife. This pattern of cognitive decline was mostly associated with reduced decoupling in the inferior parietal, posterior and mid cingulate cortices, and dlPFC, combined with reduced coupling in the SMN and visual cortices (see Table S1 – Supplementary Results).

#### 2.4.2. Semantic – Latent component 2

At a cognitive level, LC2 was mainly driven by semantic abstraction as measured by the proverb task (18.07), verbal fluency (16.45), sentence comprehension (12.49) and picture naming (7.1), after accounting for the decrease in fluid abilities (−4.19). These tasks performances peaked in midlife and remained relatively stable until very old age. This pattern of cognitive enhancement was mostly associated with increased coupling in the SMN, combined with increased decoupling in key regions including the left medial prefrontal (*area L-10r*), left middle insular (*area L-MI*), left lateral temporal (*area L-TE1*), and left posterior visual (*area L-V3CD*) areas, which appear to outline the left inferior frontal occipital fasciculus (IFOF). Similarly, increased decoupling in the right anterior cingulate (*area R-a24pr*) and right precuneus at the junction with the occipital lobe appears to outline terminations of a superior longitudinal fasciculus (SLF) branch (see Table S2 – Supplementary Results).

## 3. Discussion

Aging is marked by a simultaneous decline in cognitive control and an enrichment of semantic knowledge, which is thought to reflect a change in processing language information, and more broadly in neurocognitive mode (Spreng & Turner, 2021). Resting-state fMRI and DWI analysis have provided substantial microstructural and functional evidence for this change in midlife, with implications for processes engaging a brain-wide interaction between perceptuo-motor, control, and semantic systems that contribute to lexical production (Guichet, Banjac, et al., 2024; Guichet, Roger, et al., 2024). Yet, our studies did not account for the structural constraints on functional dynamics across the lifespan. To propose a true multimodal model of healthy aging, we proposed to examine the reorganization of structure-function-cognition relationships at rest in a population-based sample of 597 individuals aged 18-88 (Cam-CAN et al., 2014), extending the current graph signal processing framework.

Consistent with our general hypothesis, our findings highlight midlife as a critical turning point in neurocognitive reorganization characterized by distinct structure-function dynamics (Lachman, 2015; Park & Festini, 2016). Our main findings are three-fold: (1) Healthy aging recalibrates the structure-function relationship underlying integrative processing, with midlife emerging as a key transition point. Below, we elaborate on the potentially maladaptive or compensatory nature of these changes. (2) As hypothesized, the structure-function architecture supporting age-related lexical production involves both control and semantic mechanisms. (3) Structural coupling with sensorimotor processes occupied a central role for facilitating these mechanisms, helping to sustain cognitive control in younger adulthood and contributing to the development of a semantic strategy that may preserve word-finding abilities into older age.

### Age-related changes in integration: markers of decline or evidence of compensatory reorganization ?

Overall, our study replicates prior results showing that large-scale structural connectivity patterns best predict function (Behjat et al., 2023; Olsen et al., 2024; Preti & Van De Ville, 2019). We also confirm that these patterns serve as foundational structural axes for integrative processing (Sipes et al., 2024), as shown by graph-metrics in Figure 2B.

With advancing age, individuals exhibit a different structure-function architecture for integration, which may have broader implications for cognitive flexibility (Fernandez-Iriondo et al., 2025). Older adults engaged a broader repertoire of integrative harmonics to compose fMRI activity at rest. In other words, the fMRI signal is not dominated by just one or two modes but is instead supported by multiple integrative components. Despite this expanded repertoire, functional activity became increasingly independent of these integrative harmonics with age, meaning that the fMRI signal is globally more free from immediate structural constraints, likely reflecting reduced global inhibition in older adulthood (Atasoy et al., 2016; Luppi et al., 2023).

Taken together, one possible interpretation is that these effects reflect a trade-off between *increased diversity* and *decreased specificity* in the neural code governing integrative structure-function dynamics. Older adults tend to recruit additional large-scale structural connectivity patterns, yet these may not fully serve functional processes which demand spatially-specific patterns of integration, for example along specific structural axes. This interpretation gains further support when considering that the structural topology is also less supportive of moment-to-moment fMRI dynamics in older adulthood, thus also affecting highly-synchronized processes such as inhibitory control (Courtney & Hinault, 2021).

Whether this trade-off represents a maladaptation, compensation, or a mix of both as noted by McDonough et al. (2022), remains unclear. *Maladaptation* may take place in face of substantial losses of microstructural degeneration along the main axes of structural connectivity, especially those involved in cognitively demanding tasks such as during language processing (Kljajevic & Erramuzpe, 2019; Sánchez et al., 2023; Troutman et al., 2022; Yeske et al., 2021). This could lead to additional recruitment of large-scale anatomical pathways with limited little to no benefits for cognition, akin to processes of functional dedifferentiation (for a review, see Deery et al., 2023; Koen et al., 2020). In comparison, *compensation* may reframe this recruitment as structural adaptations that provide a more balanced and redundant structural topology for integration, providing robustness to integrity loss along the main structural axes, at the cost of less “precise neural code”, as noted by Johnston & Freedman (2023).

### Age-related changes in integration: evidence of a transition to a exploitative neurocognitive mode

Our study further reveals spatial differences in structure-function integration across age groups. As shown in Figure 2A, older adults showed greater integrative processing in the posterior medial prefrontal cortex (mPFC), while younger adults relied more on anterior mPFC activity. This aligns with theories of goal-directed behavior, emphasizing that the posterior PFC exploits the control resources to optimize task execution, while the anterior PFC facilitates the monitoring and redistribution of cognitive resources when managing competing goals – a crucial element to cognitive flexibility. This functional dissociation between an “exploratory” and “exploitative” drive for goal-directed behavior (Mansouri et al., 2017; Soltani & Koechlin, 2022; but see also Badre & Nee, 2018) fits well with the “exploration-to-exploitation shift” in neurocognitive mode proposed by Spreng & Turner (2021) and hypothesized in this study.

Additionally, we observed a ventro-dorsal reconfiguration in posterior parieto-occipital cortices, which echoes recent MEG findings from the same CamCAN dataset (Guichet, Harquel, et al., 2024). These findings suggested that the observed antero-posterior and ventro-dorsal axes of reconfiguration accompany a shift toward faster, alpha-dominant dynamics in older adulthood. However, the behavioral relevance of this shift with regard to structural connectivity remains inconclusive. For example, Gao et al. (2020) reported that brain-wide acceleration of neural dynamics with age negatively impacts the integration of long-term cues during working memory maintenance; while Blanco et al. (2024) suggested that alpha oscillations increases the coupling to the structure, likely enhancing inhibitory and top-down semantic control (Guichet, Harquel, et al., 2024; Klimesch, 2012; Zioga et al., 2024). Future work should explore time-resolved structure-function dynamics (Liu et al., 2022; Sadaghiani & Wirsich, 2020), especially considering the repertoire neural timescales managed in the PFC (Spitmaan et al., 2020).

### Structure-function underpinnings of cognitive control decline

In line with our hypothesis, we found both a cognitive control and semantic neurocognitive trajectory with inflection points at midlife. We discuss the former trajectory below and address the latter in the following subsection.

The cognitive control trajectory explained most of age-related changes, largely recapitulating the main effect of age alone reported in section 2.3. As noted previously, healthy aging weakened the relationship between large-scale structural axes and functional activity, but this effect was not uniform across functional networks. Higher-level networks, such as the DMN, DAN, and FPN, showed the most reduction in decoupling with age. This pattern further correlated with the onset of control deficits related to lexical production in midlife, as shown in Figure 4, reaffirming word finding as a multimodal phenomenon (Rahman et al., 2023). A closer examination of changes onsetting in midlife further revealed that reduced decoupling mostly impacted the PCC. This is remarkably similar to the age-related patterns of neural flexibility formalized in the SENECA model, suggesting that PCC functional dynamics are mostly free from immediate structural constraints until midlife and represent a key element for flexible goal-directed cognition across imaging modalities (Guichet et al., *in revision*).

In comparison, lower-level networks, such as the SMN, Visual, and Auditory networks, showed reduced coupling associated cognitive control deficits. Again, this echoes the substantial increase in the flexibility of lower-level processes in older adults reported in SENECA. This could be interpreted as a more energy-efficient, multi-sensory integration mechanism for accessing the control demands of lexical production in face of declining metabolic resources with age (Deery et al., 2024; Guichet et al., *in revision*).

Collectively, these results show that most of the structure-function architecture reconfigures within hierarchical groups of the unimodal-to-transmodal gradient during healthy aging (Margulies et al., 2016; Sydnor et al., 2021), further establishing this gradient as a foundational organizational principle of neurocognition across imaging modalities (Collins et al., 2024; Feng et al., 2024; Fotiadis et al., 2024; Monaghan et al., 2025; Paquola et al., 2019; Vázquez-Rodríguez et al., 2019) and of spatiotemporal functional growth across the human lifespan (Sun et al., 2025). On that note, core regions of the language network (as defined by Ji et al., 2019) showed both increased and reduced coupling with age, further suggesting that language is a highly-interactive domain that likely communicates with both high- and low-demand structure-function architectures.

### Structure-function underpinnings of the semantic strategy

After accounting for cognitive control decline, we identified the hypothesized semantic trajectory which grew from younger adulthood to midlife and remained stable with age. As noted in the Introduction, previous research had already identified a left subcortico-sensory white matter network supporting this strategy in middle-aged adults (Guichet, Roger, et al., 2024). Here, our study confirms that pathways converging on the SMN play a central role in this semantic strategy by increasing their coupling to the structure, likely providing a fast, reliable, and “phase-locked” response to stimuli in the environment (Preti & Van De Ville, 2019; Rossi-Pool et al., 2021) that bridges the external world with internal processes (Schwartz, 2016). Embodied cognition represents an interesting framework to understand the benefits of sensory-driven processing on semantic systems in older adults. More reliable sensorimotor input/output could enhance the fidelity “embodied internal models” that are generated by integrating these inputs with a lifetime of learned semantic representations (Fernandino & Binder, 2024; Martin, 2016).

The interaction between these two systems is further supported by our results, showing that enhanced structure-function coupling in the SMN operates in unison with greater decoupling in regions outlining the IFOF, a pathway crucial to long-term semantic representation (Krieger-Redwood et al., 2025; Ralph et al., 2017; Visser et al., 2010). In sum, this supports the idea that age-related lexico-semantic reorganization involves an interaction between the structurally-coupled sensorimotor and structurally-decoupled semantic system, likely establishing the foundations for delaying the onset of word finding difficulties in face of declining cognitive control (Baciu & Roger, 2024; Krethlow et al., 2024).

### The key role of structurally-coupled sensorimotor dynamics

As noted in the last two subsection, structurally-coupled SMN activity largely contributed to the control and semantic trajectories of age-related lexical production. It could act as a communication channel with structurally-decoupled activity. For example, it may sustain cognitive control until midlife via feedforward and feedback connections with structurally-decoupled activity in transmodal cortices, thereby predisposing individuals to the integration of novel experiences (Griffa et al., 2022). In parallel, it may contribute to establish a more semantic strategy for lexical production that can be maintained for longer in older adulthood, as described in the last subsection.

We may suggest key regions that interface both trajectories, that is showing an increased coupling up until midlife – establishing the semantic strategy – followed by reduced coupling in older adults – reflecting cognitive control decline and vulnerability to microstructural changes (Pan et al., 2024). These regions include the primary right somatosensory cortex (area 1), the bilateral cingulate regions located in the anterior inferior paracentral lobule (area 24dd) which show dense functional connectivity to sensorimotor but also other sensory modalities (Baker et al., 2018), and the left posterior middle frontal gyrus (pMFG - area 55b) which interfaces lightly and heavily myelinated cortical microarchitectures – enabling both low-level sensorimotor integration (Siman-Tov et al., 2022), and high-level integrative processing required for language production tasks (Chang et al., 2020; Donahue et al., 2018; Hazem et al., 2021). We also noted substantial implications of associative, callosal, and projection fibers transiting via these regions, underscoring their broad relevance for neurocognition.

In sum, our study complements a vast body of research showing how the SMN functional synchrony shapes brain-wide activity (Gordon et al., 2023; Kong et al., 2021), specifically in the PFC (Fine & Hayden, 2022; Goelman et al., 2024), with repercussions on language, memory, and motor control processes (Ferré et al., 2020; Zapparoli et al., 2022).

### Limitations of this study

This study has several limitations. **(1) Length of timeseries.** The CamCAN dataset provides around 8 min of resting-state recording with only 261 timepoints. Longer sampling time and related initiatives are needed to enhance robustness, statistical power and provide precision structure-function mapping (Gordon et al., 2017). **(2) Cross-sectional analysis.** Similarly, longitudinal datasets represent an important step forward to refine the proposes model of aging to individuals. **(3) Cortico-centric analysis.** Our results suggested that projection fibers may transit via key structure-function hubs during age-related reorganization. Future work may extend the present study to include thalamic and striatal projections when examining structurally-constrained fMRI activity. **(4) Inter-individual variability.** Individuals showed poor alignment of harmonics that lie in the middle of the energy spectrum – the structural modes that bridge integration and segregation. Future work is needed to assess the behavioral relevance of this structural regime as it was not specifically associated with age in this study, although it was found to be important for cognitive flexibility in a recent work (Sipes et al., 2024).

### Conclusion

Network neuroscience has significantly advanced our understanding of how structural and functional connectivity evolve during healthy neurocognitive aging. Yet, multimodal studies linking structural and functional brain organization with cognitive performance remain relatively scarce. Using a graph signal processing framework, this study analyzed resting-state functional MRI and diffusion-weighted imaging data from 600 healthy adults aged 18 to 88, drawn from the CamCAN dataset. Our results showed that control and semantic systems follow distinct age-related trajectories of structure–function reorganization, with midlife emerging as a critical inflection point. Before midlife, structurally-coupled sensorimotor integration coexists with structurally-decoupled activity in transmodal cortices, supporting cognitive control. After midlife, a shift occurs toward a sensorimotor-based semantic strategy, marked by greater structure–function coupling in primary systems and increased decoupling along long-range semantic pathways, potentially scaffolding embodied internal models for cognitive resilience. Together, we conclude that sensorimotor dynamics serve as a plausible scaffold for controlled semantic cognition throughout the lifespan, extending current neurocognitive aging models such as LARA and SENECA.

## 4. Material and Methods

### 4.1. Participants

Our dataset included 600 healthy adults (age range: 18-88; 295 males, 305 females) from the Cambridge Center for Ageing and Neuroscience Project (http://www.mrc-cbu.cam.ac.uk/: Cam-CAN et al., 2014). Details on participant recruitment are provided in Taylor et al. (2017).

This sample size was obtained by including the participants with at least 5 (out of 8) cognitive scores. Any missing score was imputed with the median value of the participant’s age decile. These 8 scores were derived from neuropsychological tasks assessing lexical production performances both directly and indirectly (see Table 1). As noted in section 4.3.3, three additional subjects were removed during structure-function analysis.

**Table 1.** Neuropsychological tasks and associated cognitive processes were considered in this study. A detailed description can be found in Supplementary Materials. LP=Language production.

| Neuropsychological test | Cognitive processes | Task type and relation to lexical production |
| --- | --- | --- |
| **Picture naming** | Language (phonological and semantic access) | LP, Direct |
| **Verbal fluency** | Language (phonological and semantic access) and EF | LP, Direct |
| **Tip-of-the-tongue situations** | Language (retrieval) and EF (monitoring) | LP, Direct |
| **Proverb comprehension** | Language (comprehension) and EF (abstraction) | Language, Indirect |
| **Sentence comprehension** | Language (syntax, semantic) | Language, Indirect |
| **Story recall** | Long-term memory | Language, Indirect |
| **Cattell task** | EF (Logical reasoning, problem-solving) | Domain-general, Indirect |
| **Hotel task** | EF (multitasking, planification, self-monitoring) | Domain-general, Indirect |

### 4.2. MR acquisition and preprocessing

As the data was sourced from Cam-CAN et al. (2014), we refer the reader to Taylor et al. (2017) for multi-shell diffusion-weighted MRI and resting-state functional data acquisition details.

#### Structural data

Diffusion MRI data were preprocessed using the *MRtrix3* software (Tournier et al., 2019) (version 3.0.4; https://www.mrtrix.org/). Preprocessing steps included denoising (Veraart et al., 2016), Gibbs artifact removal (Tustison et al., 2010), eddy current and motion correction with FSL (Andersson et al., 2003), and bias correction with ANTs N4. Images were upsampled to a voxel size of 1 mm^3^ using cubic B-spline interpolation (Raffelt et al., 2012).

Structural connectomes (SC) were generated as follow. First, fiber orientation distributions (FOD) were computed using the MSMT CSD algorithm (Dhollander et al., 2016; Jeurissen et al., 2014; Tournier et al., 2007) with group-averaged response functions calculated before upsampling to enable comparisons across subjects. Joint bias correction and intensity normalization were subsequently applied to these FOD images. Then, whole-brain tractography (iFOD2; 5 million streamlines, max. length = 250 mm, cutoff = 0.06, backtrack option) was performed for each individual while reducing biases with the SIFT2 algorithm (Smith et al., 2015). The corresponding SC was generated by counting the number of streamlines between each pair of nodes in the HCP-MMP 1.0 atlas (Glasser et al., 2016) (symmetric option, scaling: invnodevol).

#### Functional data

Resting-state fMRI data were preprocessed using the *fMRIPrep* software (https://fmriprep.org/en/stable/: Esteban et al., 2019). Preprocessing steps included motion correction, slice timing correction, susceptibility distortion correction, co-registration, and spatial normalization. T1-weighted images underwent skull stripping, tissue segmentation, and spatial normalization.

Denoised timeseries were aggregated using the 360-region HCP-MMP 1.0 atlas (Glasser et al., 2016). We used custom scripts to co-register the atlas to subject-space with ANTs (https://github.com/ANTsX/ANTsPy) and applied a confound removal strategy via Nilearn (https://nilearn.github.io/) that preserves signal continuity (Wang et al., 2024). Confounds included high-pass filtering, the full 24 motion parameters, the basic 2 white matter and CSF parameters, and a single global signal parameter. Mean framewise displacement was also extracted and used as a covariate in subsequent statistical models. Spatial smoothing and standardization was applied to the signal using a 6mm full-width-at-half-maximum (FWHM) kernel.

### 4.3. Structure-Function analysis

We performed structure-function analysis under the graph signal processing (GSP) framework. We developed custom scripts from the NiGSP python toolbox (Stefano Moia et al., 2024) (version 0.19.0; https://github.com/MIPLabCH/nigsp).

#### 4.3.1. Graphs and Spectral Graph Theory

A graph *G* is formally defined as: *G =* (*V*, *E*), where V is the set of vertices or nodes, and *E* is the set of edges, represented the connection weight between a pair of vertices (*i*, *j*) where *i*, *j* ∈ *V*. *G* can be represented with two key matrices: (1) the adjacency matrix *A*, a square matrix where elements 𝑎_𝑖,𝑗_ denote the weight of the edge (*i*, *j*), and (2) the degree matrix *D,* a diagonal matrix where elements 𝑑_𝑖,𝑖_ = ∑_𝑗_ 𝑎_𝑖,𝑗_denoting the sum of all edge weights for each node. Using A and D, we can define the graph Laplacian 𝐿 and its normalized form ℒ as:

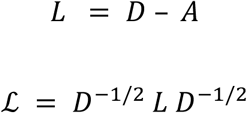

Graph Signal Processing (GSP) applies signal-processing operations to graph-structured data, assuming that the fMRI signal resides on the graph vertices *V* defined by the structural connectome *G* (Abramian et al., 2021). We only provide a brief introduction and refer to previous works for additional details (Atasoy et al., 2016; Leus et al., 2023; Lioi et al., 2021). A graph spectral representation is obtained through eigendecomposition of the normalized Laplacian: ℒ = 𝑈𝛬𝑈^𝑇^, with 𝛬 = {0 = 𝜆_1_ ≤ 𝜆_2_ . .. ≤ 𝜆_𝑚𝑎𝑥_} denoting the set of eigenvalues that define the graph Laplacian spectrum, and 𝑈 the set of Laplacian eigenvectors, also known as harmonics (Behjat et al., 2025). These harmonics carry a notion of spatial frequency: low-eigenvalue harmonics vary smoothly along the graph, whereas high-eigenvalue harmonics capture more localized variations (Lioi et al., 2021).

#### 4.3.2. Harmonic alignment procedure

Although subject-level harmonics show minor differences compared to a group-averaged decomposition (Olsen et al., 2024), the ordering of harmonics can still be ambiguous across subjects, especially in subject-specific high-frequency harmonics (Griffa et al., 2022), thus making comparisons difficult.

To resolve ordering ambiguity, we applied a procedure that aligns subject-level eigenspaces to a reference space. This reference corresponds to the decomposition of the normalized Laplacian of the group-representative SC, built using distance-dependent consensus-based thresholding (Betzel et al., 2019). The procedure consists in computing the cosine similarity between the reference and each subject’s eigenspace while ignoring polarity (i.e., absolute value). Then, the Hungarian algorithm is applied to find the optimal permutation of eigenvectors that maximizes similarity. This ensures that eigenspaces are comparable across subjects and retains the ordering of ascending eigenvalues with respect to a group-averaged SC.

Note that other matching procedures have been proposed such a bipartite matching max-flow algorithm (Sipes et al., 2024) or Procrustes alignment (Behjat et al., 2023). However, to our understanding, the former does not guarantee a perfect matching while the latter may smooth out subject-specific features in the harmonics.

#### 4.3.3. Inter-subject agreement and outliers

In line with Sipes et al. (2024), we assessed inter-subject agreement in unaligned and aligned subjects’ eigenspaces as follows: 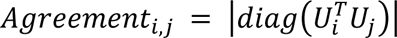. This results in an agreement value for each harmonic and *(i, j)* pair of subjects, which was averaged across pairs to highlight harmonics with the most inter-individual variability.

To identify potential outliers, we computed the Euclidean norm of the agreement between each pair of subjects. Then, we used the studentized residuals (SDR) method to examine how each subject deviates from the mean (see Figure S1 – Supplementary Results). An absolute SDR above 3 was consider as an outlier and excluded from the final sample (*N* = 597).

#### 4.3.4. Energy and diversity of structure-function relationship

Connectome harmonics serve as a transformation basis to assess how the fMRI signal aligns with structural connectivity. More specifically, the Graph Fourier Transform (GFT) projects each timepoint *t* of the fMRI signal *s* onto each harmonic *k*, thus expressing the signal into the graph spectral domain: 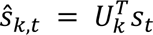.

After decomposing the fMRI signal as a linear combination of harmonics, we computed the contribution of each harmonic to each timepoint – termed energy:

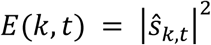

Consequently, averaging over time quantifies how much each harmonic contributes to time-averaged fMRI activity – the harmonic energy. Similarly, when averaged over harmonics, this quantifies time-specific contributions to the structural topology – the temporal energy.

In line with Sipes et al. (2024), we computed two metrics synthesizing the energy spectrum of for each subject. First, we examined harmonic diversity, reflecting the extent to which fMRI activity is composed by a wide repertoire of harmonics. Harmonic diversity was computed for each timepoint (𝐻_𝑡_) as follow and then averaged across time.

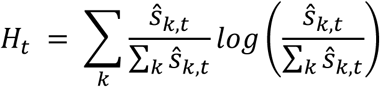

Second, we examined temporal diversity, reflecting the extent to which fMRI activity is composed by rapidly fluctuating harmonics. Temporal diversity was operationalized as the sample entropy of temporal energy. We used previously validated parameters, setting the similarity tolerance *r* = 0.2 (Richman & Moorman, 2000) and the scale parameter to match a dynamic timescale of interest in the signal with *m* = 2 (i.e., 2 TRs or about 4 s).

#### 4.3.5. Structure-Decoupling Index (SDI)

Following Preti & Van De Ville (2019), the structure-decoupling index (SDI) measures the ratio between the decoupled and coupled component of the signal. Harmonics are filtered into a set of low-frequency (coupled) and a set of high-frequency (decoupled) harmonics, with both set having roughly equal harmonic energy. The structurally-coupled 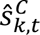 and structurally-decoupled 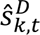 decompositions are then projected back into the graph domain using the inverse GFT: 𝑠_𝑡_ = 𝑈_𝑘_*Ŝ*_𝑘,𝑡_. Finally, the SDI is computed as the ratio of their Euclidean norms across time. A high SDI indicates high decoupling and conversely, a low SDI indicates high coupling.

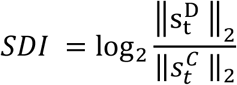

Significance was assessed by generating structurally-informed null models for each subject using the spectral randomization procedure proposed in Preti & Van De Ville (2019). Non-significant SDI values were set to 0 for further statistical analysis, indicating an equal tendency for coupling and decoupling.

### 4.4. Statistical analysis

Statistical analyses were carried out in R (version 4.3.2). All models included sex, handedness, total intracranial volume, and mean framewise displacement as covariates.

#### Main effect of age

We examined the effect of age on each metric by fitting a generalized additive model (GAM) with age as a smooth predictor (parameters: *k = 3, method = REML*). Significance thresholds were adjusted at the False Discovery Rate (FDR).

#### Structure-Function-Cognition

To examine the relationship between SDI values and lexical production performance across the lifespan, we employed Partial Least Squares (PLS) correlation analysis using the toolbox myPLS (https://github.com/MIPLabCH/myPLS) in MATLAB R2020b. PLS is a statistical technique designed to identify latent relationships between brain features (*X matrix*), here the SDI values, and cognitive (*Y matrix*) performances.

For the cognitive matrix, we included the 8 cognitive scores presented in Table 1 as well as two variables derived from piecewise polynomials. These polynomial functions allow the model to optimize the covariance between SDI values and potential nonlinear age trajectories. The first function represents an acceleration after a transition age *t,* and complementarily the second represents a leveling-off after *t*. We fixed the transition age at age 54 in line with results reported in section 2.2. This approach has been validated in previous teamwork (Guichet, Harquel, et al., 2024).

## Data and code availability

Code is made publicly available at: https://github.com/LPNC-LANG/CamCAN_Fusion

## Supplementary information

Supplementary materials : Description of the neuropsychological tests Supplementary results :

- Figure S1. Distribution of Studentized residuals to detect subject with potential outlying harmonic decomposition
- Table S1 & S2. Salient SDI changes

## CRediT author statement

Conceptualization: CG; Methodology, Formal Analysis & Data Curation: CG; Writing – Original Draft: CG, MB; Writing – Review & Editing: CG, MB, MM, SA; Funding Acquisition: MB, MM, SA; Supervision: MB, MM, SA.

## Acknowledgments

This work was supported by the ANR project ANR-15-IDEX-02 and MIAI @Grenoble Alpes (ANR-19-P3IA-0003). This project has received financial support from the CNRS through the MITI interdisciplinary programs. The Cambridge Centre for Ageing and Neuroscience (Cam-CAN) research was supported by the Biotechnology and Biological Sciences Research Council (grant number BB/H008217/1).

