## Supplementary information for "Structure-Function Dynamics in Healthy Cognitive Aging: A Graph Signal Processing Approach"

**Supplementary materials**

**Description of the 8 neuropsychological tests used to assess lexical production.** Please refer to original CamCAN articles for more information on behavioral datasets ^1,2^.

***Cattell:*** *Cattell Culture Fair Test: Complete nonverbal puzzles involving series completion, classification, matrices, and conditions ^3^.*

***Hotel Task:*** *Perform simulated tasks of a hotel manager: write customer bills, sort money, proofread adverts, sort playing cards, alphabetize a list of names. Total time must be allocated equally between tasks; there is not enough time to complete any task ^4^.* ***In this study, we applied a log transformation and subtracted it from 1 to stay consistent with a decrease as age increases (i.e., 1-log(x)).***

***Picture Naming****: Name the pictured object presented alone (baseline), then when preceded by a prime object that is phonologically related (one or two initial phonemes), semantically related (low or high relatedness), or unrelated ^5^.*

***Proverb:*** *Read and interpret three English proverbs ^6^.*

***Sentence Comprehension:*** *Listen to and judge the grammatical acceptability of partial auditory sentences that begin with an ambiguous sentence stem (e.g., “Tom noticed that landing planes…”) followed by a disambiguating continuation word (e.g., “are”) in a different voice. Ambiguity is either semantic or syntactic, with empirically determined dominant and subordinate interpretations ^7^.*

***Story Recall:*** *Listen to a short story, recall freely immediately after, then again after a delay, and finally answer recognition memory questions ^8^. Delayed recall measure used here.*

***Tip-of-the-Tongue (ToT):*** *Participants are asked to name famous faces and indicate if they know/don’t know/or have a ToT ^9^.* ***In this study, we subtracted the score from 1 to stay consistent with a decrease as age increases (i.e., 1-x).***

***Verbal Fluency:*** *Mean of letter (phonemic) fluency and animal (semantic) fluency task. For the phonemic fluency task, participants have 1 minute to generate as many words as possible beginning with the letter ‘p’. For the semantic fluency task, participants have 1 minute to generate as many words as possible in the category “animals” ^10^.*

**Supplementary results**

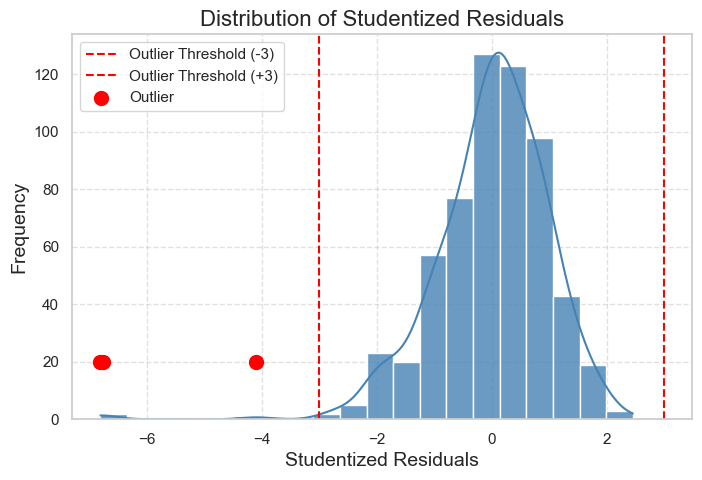

**Figure S1. Distribution of Studentized residuals to detect subject with potential outlying harmonic decomposition**

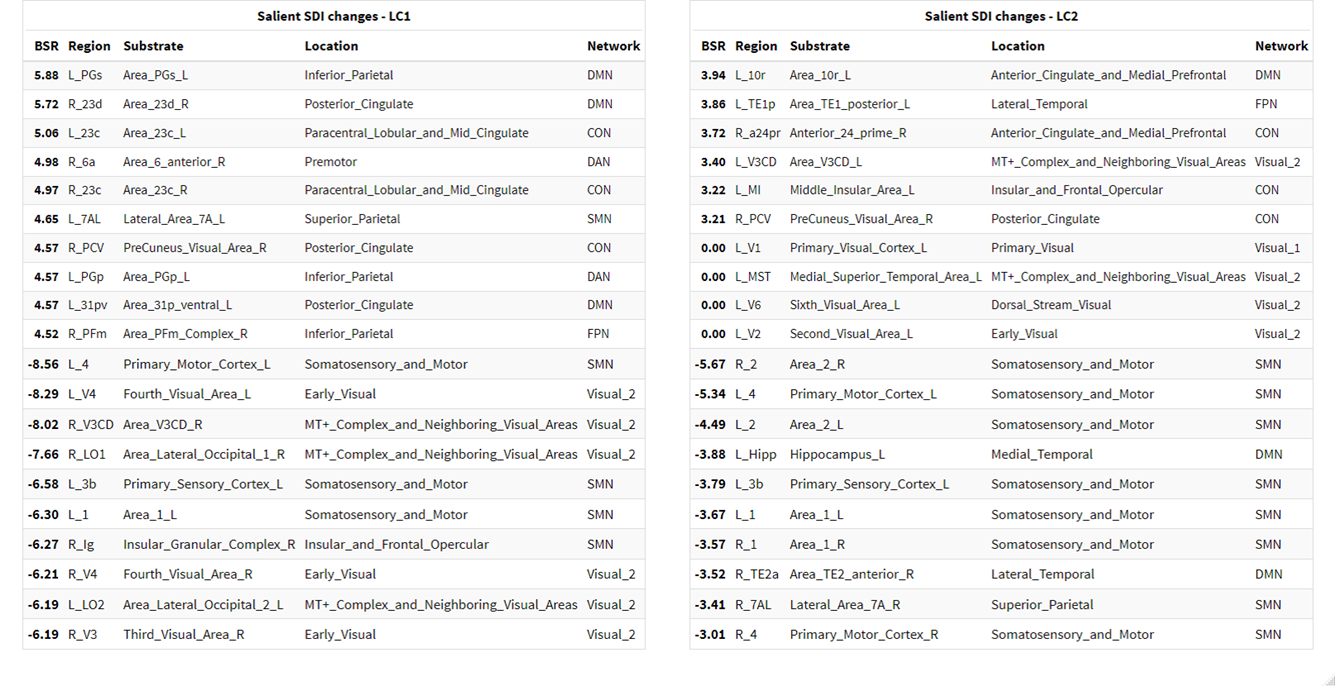

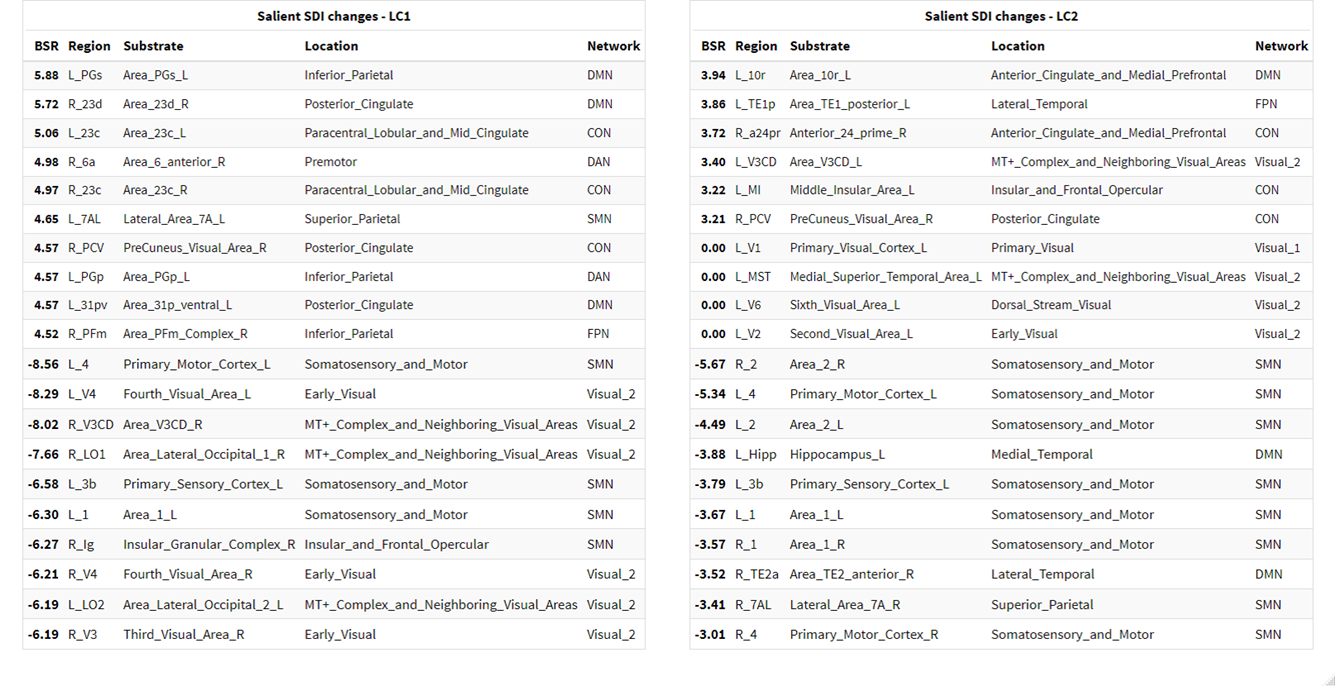

**Table S1 & S2. Salient SDI changes.** Only the most salient regions are listed here. Please refer to Figure 3 in the manuscript for a thorough depiction on the brain surface
